# Herpes Simplex Virus Glycoprotein C Regulates Low pH Entry

**DOI:** 10.1101/858472

**Authors:** Tri Komala Sari, Katrina A. Gianopulos, Darin J. Weed, Seth M. Schneider, Suzanne M. Pritchard, Anthony V. Nicola

**Affiliations:** Department of Veterinary Microbiology and Pathology, College of Veterinary Medicine, Washington State University, Pullman, WA, USA; Protein Biotechnology Graduate Training Program, College of Veterinary Medicine, Washington State University, Pullman, WA, USA

**Keywords:** herpesviruses, herpes simplex virus, viral entry, viral glycoproteins

## Abstract

Herpes simplex viruses (HSVs) cause significant morbidity and mortality in humans worldwide. Herpesviruses mediate entry by a multi-component, virus-encoded machinery. Herpesviruses enter cells by endosomal low pH and pH-neutral mechanisms in a cell-specific manner. HSV mediates cell entry via envelope glycoproteins gB, gD, and the heterodimer gH/gL regardless of pH or endocytosis requirements. HSV envelope proteins that function selectively in a given entry pathway have been elusive. Here we demonstrate that gC regulates cell entry and infection by a low pH pathway. Conformational changes in the core herpesviral fusogen gB are critical for membrane fusion. The presence of gC conferred a higher pH threshold to acid-induced antigenic changes in gB. Thus, gC may selectively facilitate low pH entry by regulating conformational changes in the fusion protein gB. We propose that gC modulates the HSV fusion machinery during entry into pathophysiologically relevant cells, such as human epidermal keratinocytes.

**Importance:** Herpesviruses are ubiquitous pathogens that cause lifelong latent infections and are characterized by multiple entry pathways. We propose that herpes simplex virus (HSV) gC plays a selective role in modulating HSV entry by a low pH pathway, such as into epithelial cells. gC facilitates conformational change of the main fusogen gB, a class III fusion protein. We propose a model whereby gC functions with gB, gD, and gH/gL to allow low pH entry. In the absence of gC, HSV entry occurs at a lower pH, coincident with trafficking to a lower pH compartment where gB changes occur at more acidic pHs. This study identifies a new function for gC and provides novel insight into the complex mechanism of HSV entry and fusion.

## Introduction

Herpesviruses contain multi-component fusion complexes and commandeer diverse entry pathways to enter target cells (1–5). Intracellular low pH facilitates entry of several herpesviruses in a cell-specific manner. The prototype alphaherpesvirus, herpes simplex virus, utilizes an acidic endosomal pathway to enter epithelial cells and a pH-independent, direct penetration pathway to enter neuronal cells (6, 7). The cellular triggers of herpesvirus entry, including intracompartmental pH, remain incompletely understood. HSV entry requires a host cell receptor that binds to viral glycoprotein D, such as HVEM or nectin-1 (8–10), but additional virus-host interactions are likely critical for entry. HSV particles contain at least 12 different virus-encoded envelope proteins. HSV entry into all cells requires gB, gD, and gH/gL. However, the majority of the remaining viral envelope proteins are not thought to be required for entry via either low pH or pH-neutral routes (11). Envelope proteins specific for a given HSV entry pathway have not been identified.

Glycoprotein B is conserved among herpesviruses and is a member of the class III fusion protein family. Unlike other class III fusion proteins such as VSV G and baculovirus gp64, herpesviral gB alone is not sufficient for fusion and requires additional viral proteins, most commonly gH/gL (12). Activation and regulation of the fusion function of gB are incompletely understood. The gH/gL complex is thought to positively regulate gB (13–15). HSV-1 gB undergoes conformational changes during fusion and entry (16, 17). Low pH specifically induces reversible changes in gB domains I and V, which comprise a functional region containing hydrophobic fusion loops (16). Acid-triggered changes in specific gB epitopes correlate with fusion activity: (i) HSV particles entering by endocytosis have reduced reactivity with gB domain I antibody H126, and elevation of endosomal pH blocks this change (16), (ii) irreversible acid-triggered changes in the H126 epitope coincide with irreversible acid-inactivation of HSV fusion and entry (18), and (iii) a hyperfusogenic form of gB has reduced reactivity with domain I and domain V antibodies, similar to low pH-treated gB (19). Thus, the acidic milieu of endosomes may serve as a host cell trigger of gB function.

HSV-1 gC, a 511 amino acid, type I integral membrane glycoprotein, mediates HSV-1 attachment to host cell surface glycosaminoglycans. This interaction is not essential for HSV entry (20–22). Here we report that gC regulates low pH viral entry independent of its known role in cell attachment. We demonstrate that gC facilitates low-pH-induced antigenic changes in gB and that gC enhances the ability of HSV to enter and infect cells by a low-pH pathway. The results are consistent with a model: in the absence of gC, HSV entry occurs at a lower pH, coincident with trafficking to a lower pH compartment where gB changes occur at more acidic pHs. We propose that gC modulates HSV entry mediated by gB, gD and gH/gL into physiologically relevant cell types, such as human keratinocytes.

## Results

### HSV entry requires intracellular low pH in a cell-type dependent manner

Herpesviruses commandeer acidic endosomal pathways to enter physiologically relevant host cells, a concept that was first demonstrated for HSV (1, 6, 7, 23–28). HSV is proposed to utilize distinct cellular routes to infect its main target cells in the human host. HSV enters epithelial cells, the site of lytic replication by a low pH mechanism, and neurons, the site of latent infection, via a pH neutral one. HSV entry into human keratinocyte cell lines HaCaT and HEKa and into model CHO-HVEM cells is inhibited by lysosomotropic agents that elevate the normally low pH of endosomes (Table 1). In contrast, entry into human neural cells IMR-32 and SK-N-SH and into model Vero cells are not blocked by lysosomotropic agents (Table 1).

**TABLE 1.** Cells used in this study and the role of low pH in HSV entry

| Cell | Type | Low pH-dependent entry of HSV<sup>a</sup> | Reference |
| --- | --- | --- | --- |
| CHO-HVEM | Chinese hamster ovary cell expressing HVEM | yes | (7) |
| HaCaT | Human epidermal keratinocyte | yes | (6) |
| HEKa | Primary adult human epidermal keratinocyte | yes | (6);This study |
| IMR-32 | Human neuroblastoma | no | (6) |
| SK-N-SH | Human neuroblastoma | no | (6) |
| Vero | African green monkey kidney | no | (35) |

<sup>a</sup>Inhibited by ammonium chloride, bafilomycin A1, or monensin

### HSV-1 gC facilitates entry and infectivity of cells that support a low pH entry mechanism

HSV envelope glycoproteins gB, gD, and gH/gL are required for entry regardless of a role for intracellular pH in a particular cell type (23, 29–32). A survey of seven additional viral envelope proteins indicated that HSV gE, gG, gI, gJ, gM, UL45, or Us9 are dispensable for entry by either low pH or pH-neutral pathways (11). In this study, we probed the role of gC in low pH entry by employing an HSV-1 KOS strain with the gC gene deleted (HSV-1 ΔgC2-3 or ΔgC) and a repaired version of this virus containing the wildtype gC gene (HSV-1 gC2-3R or gCR) (33).

HSV-1 gC is widely recognized to initiate the viral entry process by attaching to host cell surface glycosaminoglycans, principally heparin sulfate proteoglycans (22, 34). When gC-negative HSV-1 is added to cells, from 20 to 60 min p.i. there is a delay in entry relative to wild type HSV-1 (20). However, by 90 min p.i., penetration of wild type and gC-null viruses are indistinguishable. HSV-1 lacking gC has a 1 log defect in infectivity (20). Thus, while gC is dispensable in cell culture, it is important for the viral replicative cycle.

The contribution of gC to entry of HSV-1 by a low pH pathway was evaluated. HSV entry into CHO-receptor cells and human keratinocytes proceeds via a low pH endocytic pathway and is well-characterized (Table 1) (6, 7). The efficiency of ΔgC infection of CHO-HVEM cells and primary human keratinocytes (HEKa) was compared to Vero and IMR-32 cells, which support pH-neutral entry via penetration at the plasma membrane (35, 36). To control for attachment, virus was first added to cells at 4°C for 1 hr. Following a shift to 37°C for 6 h, the percentage of viral antigen-positive cells was quantitated. The efficiency of HSV-1gCR entry into each of the four cell types was similar under the conditions tested (Fig. 1A). In contrast, entry of HSV-1 ΔgC into CHO-HVEM and HEKa cells was ∼50% and 35% less efficient than into Vero and IMR-32 cells, respectively (Fig. 1B). This suggests that gC contributes to low pH entry of HSV.

**Fig. 1.**
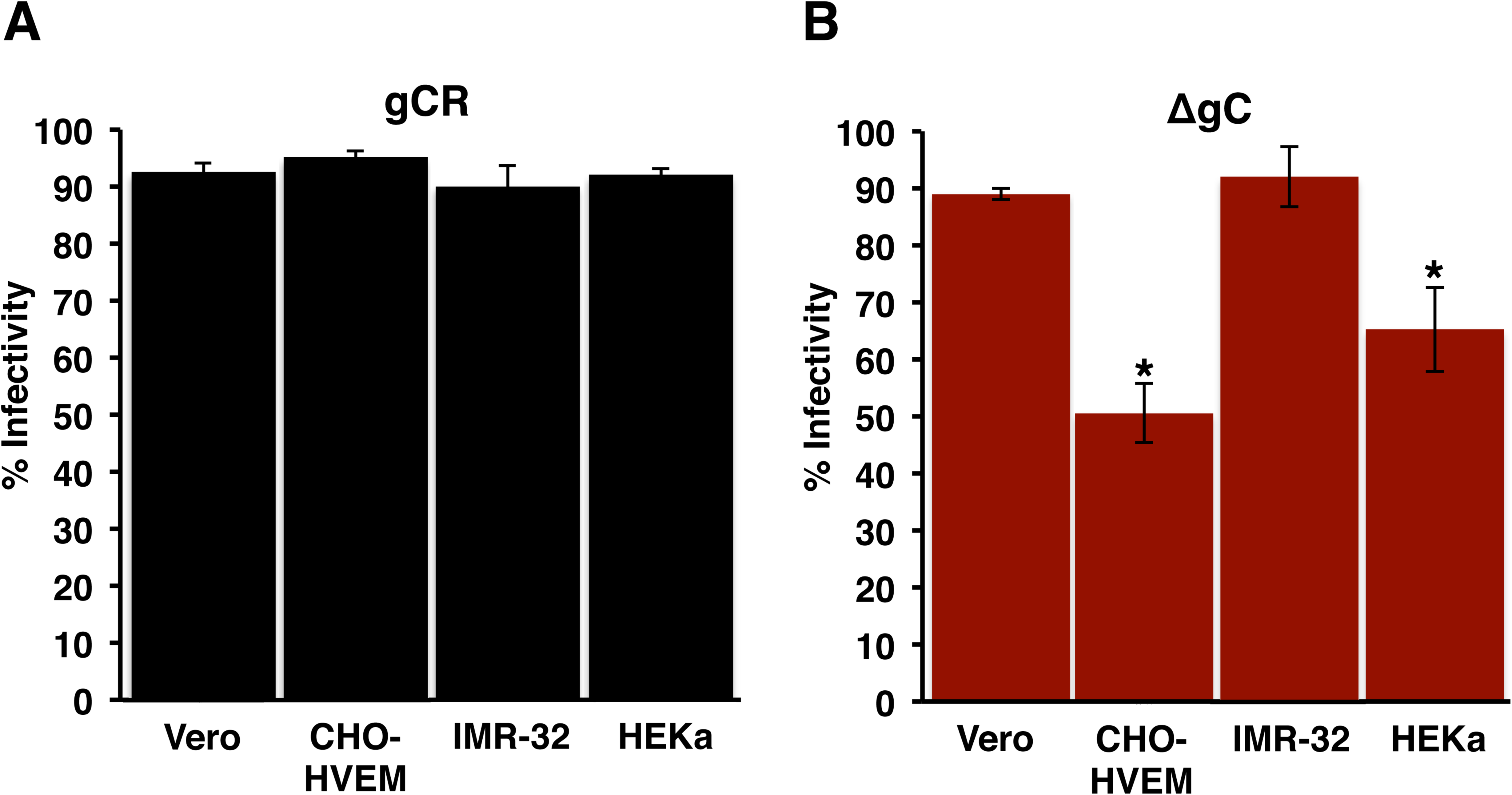
Reduced low pH entry and infectivity of HSV-1 lacking gC. Equivalent inocula of HSV-1 gCR (A) or ΔgC (B) was bound to Vero, CHO-HVEM, IMR-32, or HEKa cells for 1 h at 4°C. Following shift to 37°C for 6 hr, infected cells (MOI < 1) were quantitated by immunofluorescence. Infectivity is measured as percent HSV antigen-positive cells of ∼ 500 total cells. Student’s *t* test (*, *P <* 0.05).

### gC contributes to HSV plating efficiency on cells that support a low pH entry pathway

To confirm and extend this conclusion using an alternate approach, the plating efficiency of HSV-1 ΔgC on different human cell lines was tested. The neuroblastoma SK-N-SH line supports pH-neutral entry of HSV, and HaCaT epidermal keratinocytes support low pH entry (Table 1) (6). Identical preparations of HSV-1 ΔgC or gCR were titered. gCR had a similar plating efficiency on SK-N-SH and HaCaT cells (Fig. 2A). ΔgC had ∼ 1 log lower plating efficiency on HaCaT cells than on SK-N-SH cells (P < 0.01) (Fig. 2B). This suggests that gC is specifically important for HSV infectivity of HaCaT cells, which support a low pH entry pathway.

**Fig. 2.**
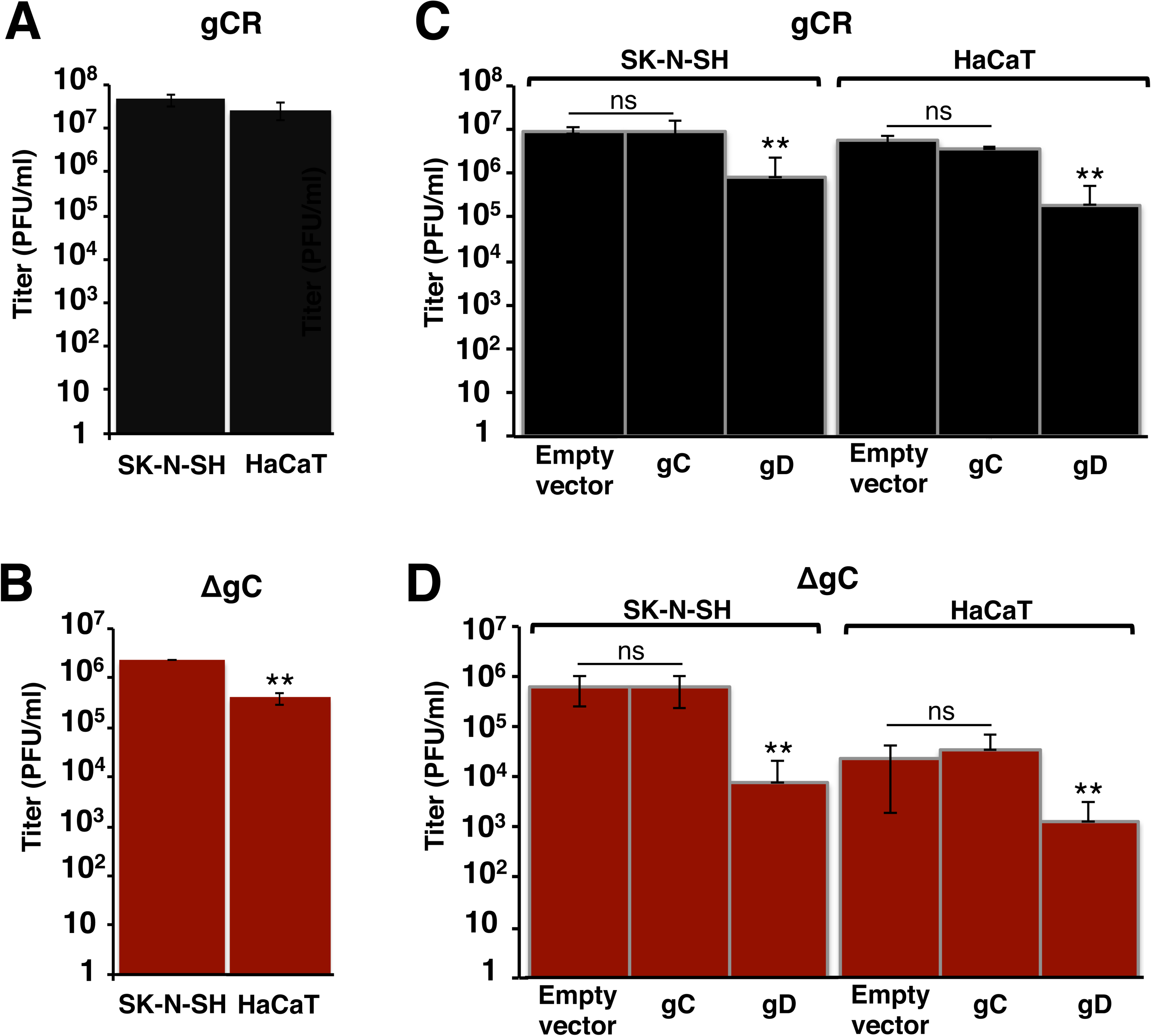
Efficiency of gC-negative HSV-1 infection of human cells that support pH-neutral or low pH entry. Equivalent inocula of HSV-1 gCR (A) or ΔgC (B) were added to SK-N-SH or HaCaT cells at 37°C. At 18-24 h p.i., titers were determined by plaque assay. (C, D) Attempt to restore infectivity of HSV-1 lacking gC by providing gC in the cell. Equivalent inocula of HSV-1 gCR (C) or ΔgC (D) were added to SK-N-SH or HaCaT cells transfected with empty vector, gC, or gD plasmids at 37°C. At 18-24 h p.i., titers were determined by plaque assay. Student’s *t* test (**, *P* < 0.01).

To investigate a potential mechanism of gC’s involvement in low pH entry, we determined whether ectopic expression of gC restored the infectivity defect of ΔgC in HaCaT cells. SK-N-SH or HaCaT cells were transfected with gC or gD plasmids, and then the plating efficiency of gCR or ΔgC was determined. Cellular expression of gC had no effect on the infectivity of gCR or ΔgC (Fig. 2C, D). Cell-expressed gD reduces HSV entry and infectivity by competing with virion gD for receptor binding (37). As expected, ectopic expression of gD in either of the cell types reduced infectivity of both gCR and ΔgC (Fig. 2C, D). These results suggest that gC functions in low pH entry by playing a role other than receptor-binding.

### gC has no detectable effect on the protein composition of HSV particles

To address the possibility that deletion of gC might affect the incorporation of gB or other viral proteins into the HSV particle, the protein composition of ΔgC was compared to gCR. The absence of gC from virions did not measurably alter the protein composition of particles as measured by SDS-PAGE and protein staining (Fig. S1A). Envelope proteins gB, gD, gE, and gH were detected to an equivalent extent in both viruses by Western blot (Fig S1B). As expected, gC was not detected in ΔgC virions. The rescuant gCR contains an equivalent amount of gC as the wild type KOS parent (33) (data not shown). These results are consistent with gC playing a specific role in low pH entry and infectivity of HSV.

### Effect of ammonium chloride on low pH entry of ΔgC HSV

The quartet of gB, gD, and gH/gL are essential for pH-neutral and low pH entry. gC is dispensable for pH-neutral entry (38). To determine whether gC was essential for low pH-dependent entry of HSV, we tested the effect of ammonium chloride treatment of CHO-HVEM, HaCaT, and HEKa cells on ΔgC entry using a reporter assay for entry. Ammonium chloride blocks wild type HSV entry into these cells (Table 1). Ammonium chloride inhibited ΔgC entry into CHO-HVEM, HaCaT, and HEKa cells in a concentration-dependent manner (Fig. 3D-F). gCR was similarly inhibited. Together the results suggest that gC contributes to low pH entry (Figs. 1, 2), but is not by itself a viral determinant of selection of the low pH pathway (Fig. 3).

**Fig. 3.**
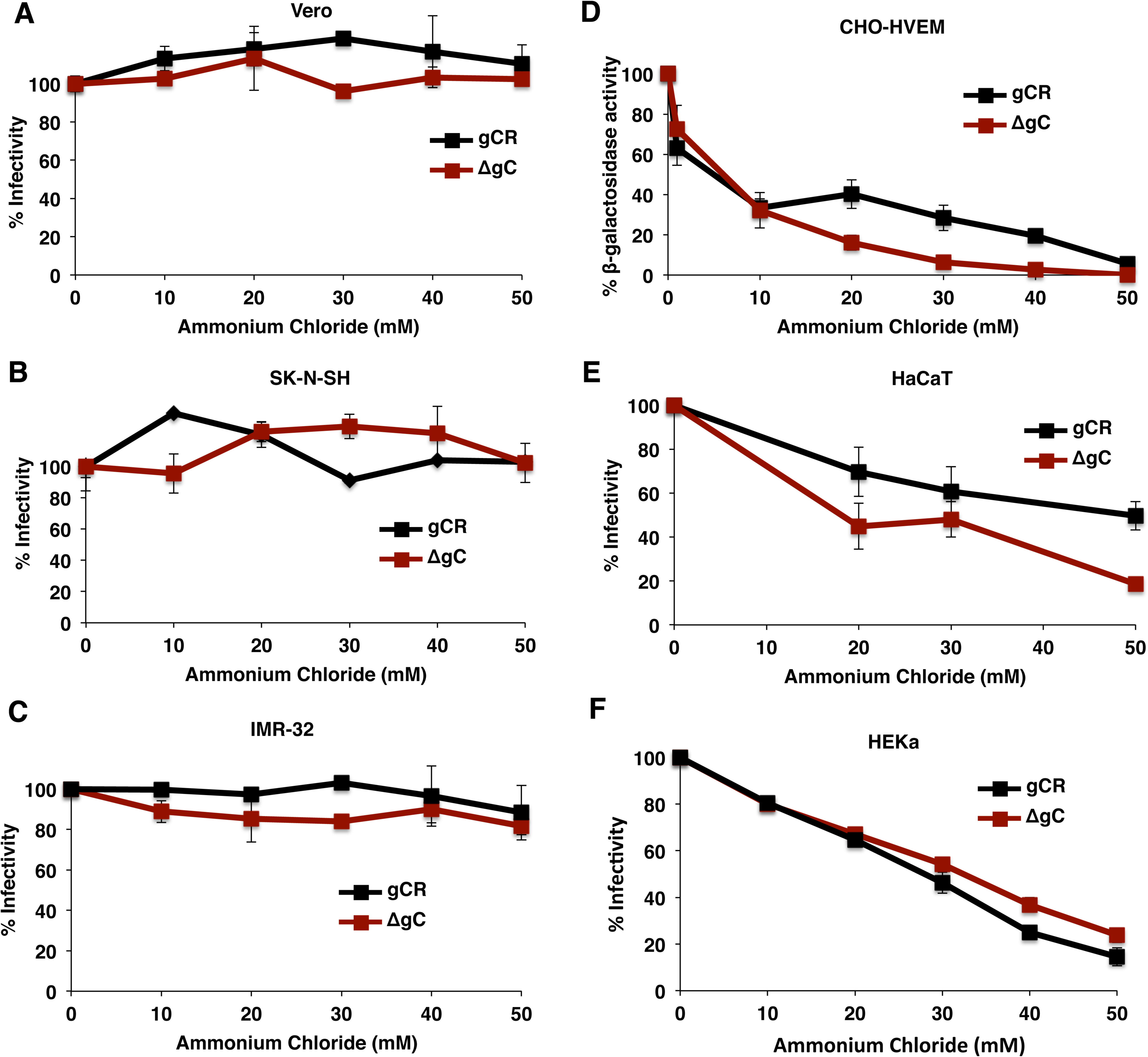
HSV-1 ΔgC enters cells via a low pH-dependent pathway. Vero (A), SK-N-SH (B), IMR-32 (C), HaCaT (E), or HEKa (F) cells were treated with ammonium chloride for 1 h at 37°C. Cells were infected with 100 PFU of ΔgC or gCR for 6 h in the continued presence of drug. Normal medium was added, and at a total of 22 h p.i., infectivity was determined by plaque assay. The infectivity of no-drug samples was set to 100%. CHO-HVEM cells (D) were treated with ammonium chloride for 20 min at 37°C. HSV-1gCR or ΔgC was added to cells (MOI of 5) at 37°C in the continued presence of agent. At 6 h p.i., entry was measured as a percent of beta-galactosidase activity obtained in the absence of ammonium chloride.

Ammonium chloride had little to no inhibitory effect on gCR entry into Vero, SK-N-SH or IMR-32 cells (Fig. 3A-C), which is consistent with pH-neutral entry of wild type HSV in these cells (Table 1). Entry of ΔgC into Vero, SK-N-SH or IMR-32 cells was similarly unaffected by ammonium chloride (Fig. 3A-C), consistent with the notion that gC is dispensable for pH-neutral entry of HSV (38).

### gC does not contribute to viral attachment under the conditions tested

We assessed the role of gC in HSV-1 attachment to the cell types used in this study. HSV-1 ΔgC or gCR was added to Vero, CHO-HVEM, SK-N-SH, HaCaT, IMR-32, or HEKa cells on ice for 1 hr at 4°C. Cell-attached HSV-1 was quantitated by qPCR. ΔgC attached to all cells in a manner similar to gCR (Fig. 4A, B). CHOpgs745 cells lack a gene required for heparan sulfate biosynthesis. Both viruses exhibited defective attachment to control CHOpgs745 cells (Fig. 4A, B). These results suggest that the altered entry and infectivity phenotypes of HSV-1 ΔgC cannot be explained by a defect in HSV-1 attachment.

**Fig. 4.**
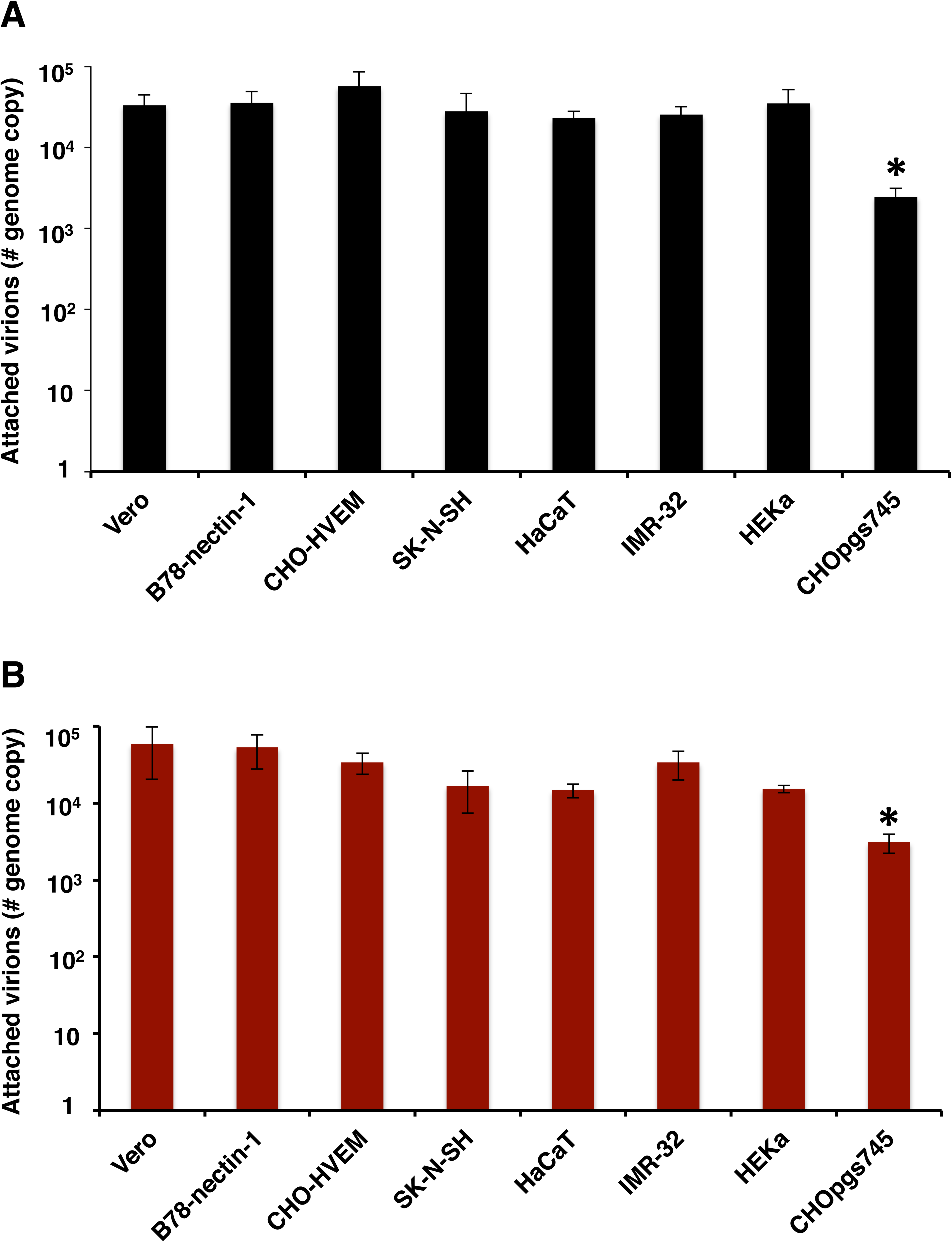
HSV-1 ΔgC attachment to cells. (A, B) 10^6^ genome copies of HSV-1 gCR (A) or ΔgC (B) were added to the indicated cell monolayers on ice at 4°C for 1 hr. Following PBS washes, cell-associated HSV-1 was quantitated by qPCR. CHOpgs745 cells lack heparan sulfate receptors for HSV attachment and serve as controls. One way-ANOVA (*, *P* < 0.01).

### gC drives the kinetics of viral penetration from acidic vesicles following intracellular transport of endocytosed HSV

To probe further the role of gC in low pH entry, we monitored the kinetics of intracellular transport of HSV-1 (23). Virus was attached to cells at 4^°^C for 1 hr. Cultures were then shifted to 37^°^C, and at different times post-infection (p.i.), the remaining extracellular virions were inactivated, and cells were lysed. The detection of infectious HSV in cell lysates reflects enveloped HSV present within cellular endocytic compartments. As expected, HSV uptake into vesicles was rapid. At 10 min p.i. in CHO-HVEM cells, there was a peak of intracellular, infectious gCR virus (Fig. 5A). Following endocytic uptake, HSV fuses rapidly with the endosomal membrane and releases its capsid into the cytosol (23). This was reflected in the sharp decrease in infectious enveloped gCR recovered by 20 min p.i. (Fig. 5A). Interestingly, ∼50% of infectious, intracellular ΔgC was recovered as late as 40 min p.i., suggesting a delay in HSV fusion with endocytic compartments in the absence of gC (Fig. 5A). Analysis of HSV-1 gCR and ΔgC trafficking in HEKa cells yielded results similar to CHO-HVEM cells (Fig. 5B). For ΔgC entry into the primary human keratinocytes, there was a ∼40 min lag in the intracellular transport and exit of HSV relative to gCR (Fig. 5B). The results from Figure 5 suggest an important role for gC in the first ∼ 20 min of wild type infection. The absence of gC appears to be overcome by ∼ 60-120 min p.i., perhaps reflecting why gC is not absolutely essential for low pH entry when longer-term assays are employed. Together, the results suggest that gC mediates rapid intracellular transport of enveloped HSV or may aid in rapid exit of HSV from acidic intracellular vesicles.

**Fig. 5.**
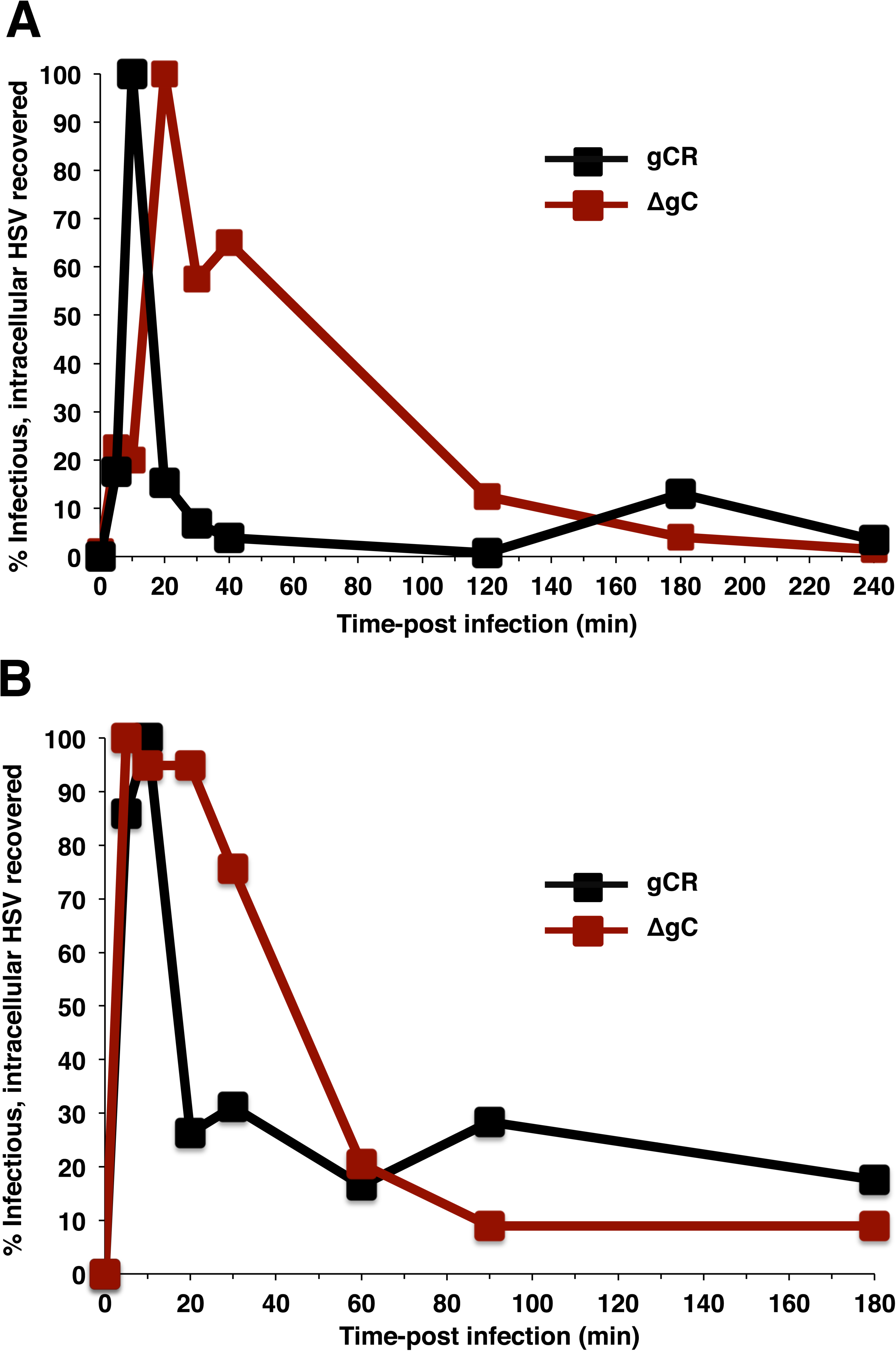
gC contributes to rapid HSV penetration following endocytosis. HSV-1 ΔgC or gCR was bound to CHO-HVEM (A) or HEKa (B) cells for 1 h at 4°C (MOI of 8). Following shift to 37°C, at the indicated times p.i., extracellular virions were inactivated, cell lysates were titered as an indication of infectious, enveloped, intracellular particles. This allows monitoring of viral trafficking and penetration over time. Peak recovery was set to 100%.

### gC positively regulates low pH-induced antigenic changes in gB

To further delineate the mechanism underlying gC’s role in low pH entry, the effect of gC on low pH-triggered antigenic changes in gB was assessed. The prefusion conformation of gB in the virion envelope undergoes low pH-triggered changes in gB domains I and V (16). These changes are at least partially reversible (16, 39, 40) and are thought to be important for membrane fusion (16, 18, 19). Domain I of gB contains internal hydrophobic fusion loops that are critical for membrane fusion (41, 42). ΔgC or gCR virions were treated with a range of mildly acidic pHs (5.0 to 7.3) and blotted immediately to nitrocellulose membrane. Blots were probed at neutral pH with nine monoclonal antibodies (MAbs) to distinct epitopes in gB, and antibody reactivity was detected and quantitated (Fig. 6). A single representative dot blot experiment for one antibody from each of six gB structural domain are shown in Figure S2.

**Fig. 6.**
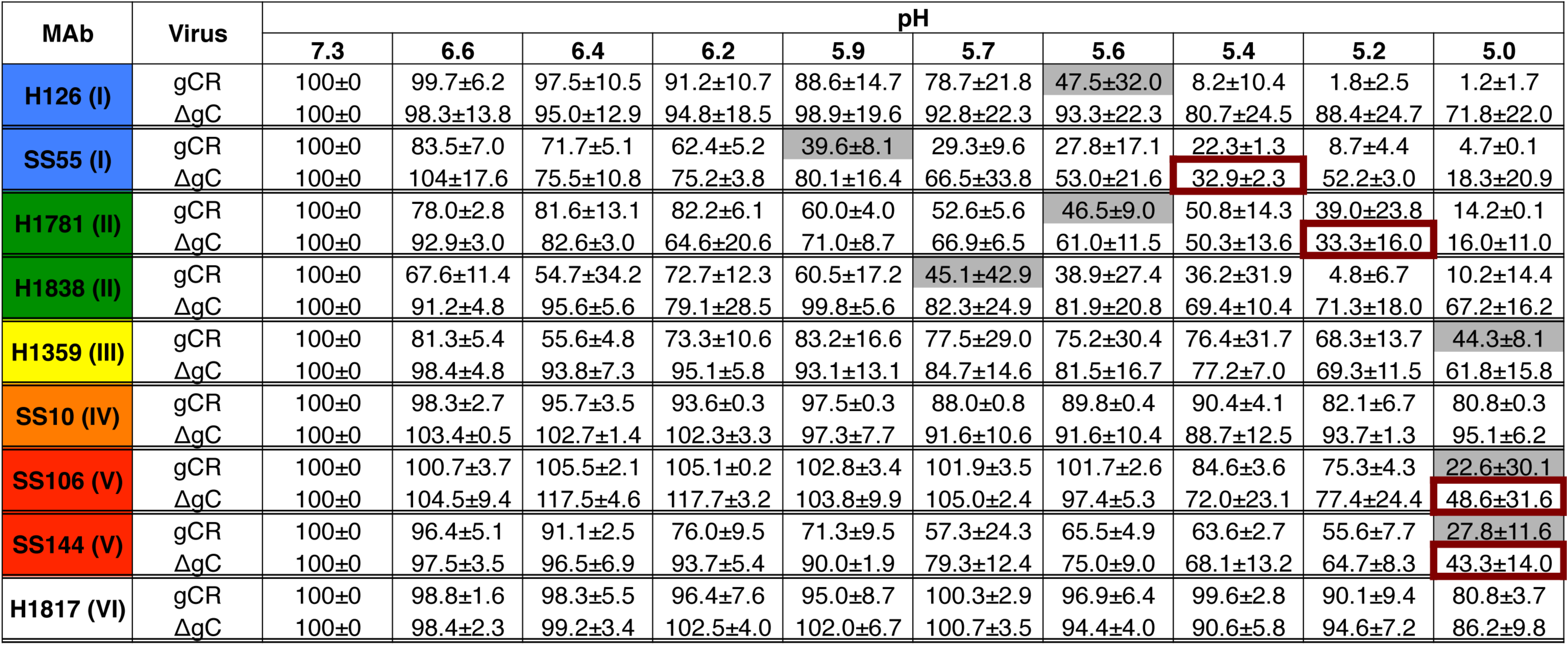
HSV-1 gC influences the pH of conformational changes in the fusion protein gB. Extracellular preparations of HSV-1 gCR or ΔgC (∼ 10^7^ genome copy numbers) were treated with pHs ranging from 7.3 to 5.0 and blotted directly to nitrocellulose. Blots were probed with gB MAbs at neutral pH. Antibody reactivity was quantitated with ImageJ. Reactivity of pH 7.3 samples were set to 100%. The pH treatment that reduced reactivity by > 50% is shaded grey (gCR) or outlined in red (ΔgC). After each MAb, the gB domain (I to VI) containing each MAb epitope is indicated (see Fig. S4).

As a reference for comparing ΔgC to gCR, the pH treatment that reduced MAb reactivity by > 50% is indicated (Fig. 6). Domain I MAb H126 had reduced reactivity with gB from gCR that had been acid-treated, but interestingly exhibited little reduction with gB from ΔgC that had been similarly treated (Fig. 6). Using 50% reactivity as a reference point, in the absence of gC, the pH at which changes in the accessibility of the H126 gB epitope occurred was reduced by at least 0.6 pH units (Fig. 6). Similar results were obtained with SS55, another MAb to gB Domain I. For SS55, the pH of gB conformational change in ΔgC was reduced by ∼ 0.5 pH units relative to gCR (Fig. 6). We have not previously examined the effect of low pH on gB Domain II. Interestingly, MAbs to domain II, H1781 and H1838, had reduced binding to gCR virions that had been treated with mildly acidic pH, suggesting that gB domain II undergoes pH-triggered conformational change (Fig. 6 and S2). The pH of antigenic change in both of the Domain II epitopes tested was decreased in the absence of gC (Fig. 6). H1838 reactivity with gB in the ΔgC virus was particularly resistant to low pH; treatment with pH 5.0, the lowest tested, still resulted in >50% reactivity, suggesting that gC alters the pH of antigenic change in the H1838 epitope of gB by > 0.7 pH units (Fig. 6). Domain III MAb H1359 had > 50% reduced reactivity only with gB from gCR that had been treated with the most acidic condition tested, pH 5.0. The H1359 epitope in HSV-1 ΔgC was more resistant to pH 5.0-treatment relative to gCR.

Acidic pH triggers specific, not global, changes in gB conformation, as low pH does not induce changes in the SS10 (domain IV) or H1817 (domain VI) epitopes of gB (16). The SS10 and H1817 epitopes in both gCR and ΔgC viruses were not altered by pH (Fig. 6). In contrast, gB Domain V from wild type HSV is known to undergo pH-induced conformational change. Following pH 5.0 treatment, Domain V MAbs SS106 and SS144 had a > 50% reduction in reactivity with gB from both gCR and ΔgC, suggesting that gC has an effect on antigenic change in gB domain V (Fig. 6).

Control deletion of a viral envelope glycoprotein gene other than gC did not alter the pH of gB antigenic change of a representative epitope (Fig. S3). The gB H126 epitope (domain I) in HSV that lacks gE (HSV-1 F-gE/GFP), underwent similar pH-induced changes relative to gB from wild type virus strain F. As a control, the H1817 (domain VI) epitope was unaffected by low pH regardless of the presence of gE (Fig. S3). Together, the results suggest that HSV-1 gC specifically increases the pH threshold of gB conformational change, particularly in Domains I and II. This is consistent with gC facilitating low pH entry and infectivity of HSV (Fig. 1, 2, 5), possibly at the level of fusion with an endosomal membrane. Cargo transiting the host lysosome-terminal endocytosis pathway is subjected to decreasing pH, from ∼ 6.5 to 4.5. HSV co-localizes with endocytosis markers, but the specific fusion compartment has not been identified (6, 43). Intracellular transport of gC-negative HSV may be delayed because transit to a lower pH compartment is necessary for fusion-associated changes in gB.

### The pH threshold of reversibility of conformational changes in gB

Reversibility of pH-triggered changes is a hallmark of gB and other class III fusion proteins (16, 44, 45). Pre-fusion and post-fusion forms of class III proteins are proposed to exist in a pH equilibrium that is shifted to the post-fusion state by acidic pH (46). During HSV egress, reversibility may allow gB on progeny virions to avoid nonproductive activation during transport through low-pH secretory vesicles. The pH threshold of initiation of gB conformational change is ∼ 6.2 to 6.4 (16). The pH at which acid-treated gB reverts to a pH-neutral conformation is not known. To determine this, virions were treated with pH 5 or maintained at pH 7.3 for 10 min at 37°C. The pH 5-treated samples were increased to different target pHs for 10 min at 37°C to determine the pH at which reversibility occurs. Samples were then blotted to membrane and probed with representative MAbs to gB (Fig. 7). Upon treatment with a low pH of 5, there was a reduced reactivity of MAb H126 (domain I) (Fig. 7A), H1781 (domain II) (Fig. 7B), and SS144 (domain V) (Fig. 7C), but not H1817 (Domain VI) (Fig. 7D). Consistent with Figure 6, the reduction of gB antibody reactivity with ΔgC was less pronounced. Upon increasing the pH of gCR from 5.0 to 5.6, there was a partial restoration of gB antibody reactivity. Increasing to pH 6.2-6.6 resulted in restoration of >80% of the reactivity measured at pH 7.3. This suggests that the pH threshold of reversibility of HSV gB conformational change is between pH 5.0 and 5.6. gB appears to be conformationally labile in this pH range.

**Fig. 7.**
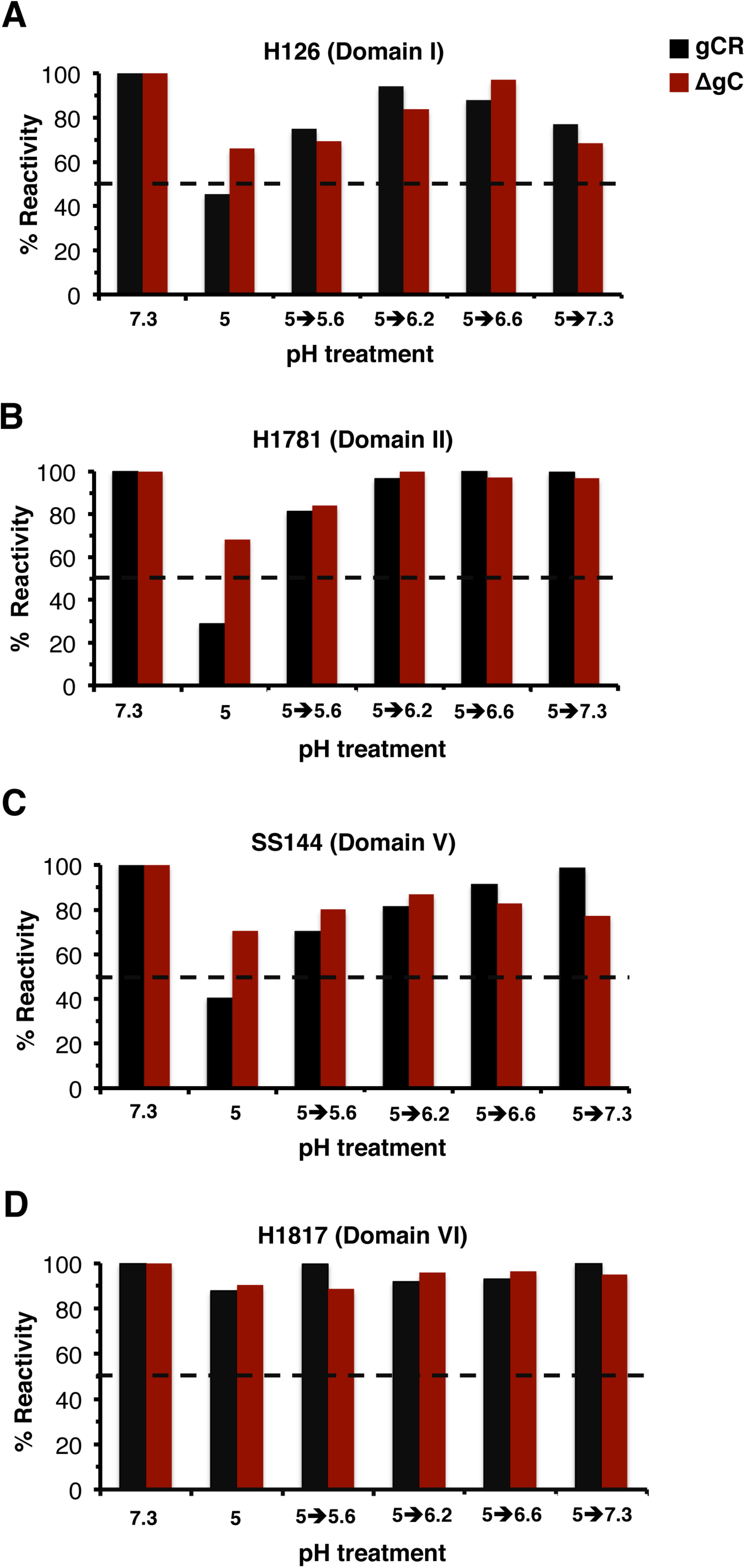
The pH threshold of reversibility of gB conformational changes. HSV-1ΔgC or gCR was treated with pH 7.3 or 5.0 and incubated for 10 min. The pHs of the pH 5-treated samples were increased to the indicated pHs for an additional 10 min. Samples were directly blotted to nitrocellulose and probed at neutral pH with gB MAb H126 (A), H1781 (B), SS144 (C), or H1817 (D). MAb reactivity was quantitated, with the pH 7.3-treated sample set to 100%.

To assess the influence of gC on the reversibility of gB conformational changes, ΔgC virions were tested for the reversibility of pH-triggered changes in the H126, H1781, SS144, and H1817 epitopes of gB. The results suggest that gC had little influence on the reversibility of gB conformational changes as measured here (Fig. 7A-C), rather gC has greater impact on the initial low pH-induced antigenic changes in gB (Fig. 6 and S2).

### Mildly acidic pH changes in gB oligomeric conformation are independent of gC

An independent indicator of pH-induced alterations in gB is a shift to a lower density gB oligomer in response to acid treatment (16). The role of gC in gB oligomeric changes was tested with MAb DL16, which recognizes an oligomer-specific epitope within gB domain V (47).

When either ΔgC or gCR were pretreated with low pH, DL16 reactivity was similarly reduced (Fig. 8A), distinguishing the DL16 epitope on gB from the other acid-sensitive epitopes (Fig. 6). This result signifies acid-triggered change in the oligomeric conformation of gB (Fig. 8A) regardless of gC’s presence. This outcome was confirmed by an independent measure of gB oligomeric conformation. When HSV is first treated with low pH and then subjected to 1% SDS and native PAGE, the slower migrating, higher molecular weight (HMW) species of gB oligomer disappears, suggesting a change in gB oligomeric conformation (16). Using this approach, the HMW gB oligomer from ΔgC disappeared in a manner similar to gB from the control rescuant virus gCR (Fig. 8B). Although gC regulates low-pH induced antigenic changes in gB (Fig. 6), it does not appear to affect acid-triggered alterations in the gB oligomer. This is consistent with the notion that a similar low pH triggers both antigenic and oligomeric alterations in gB, but these changes are experimentally separable and not identical (18).

**Fig. 8.**
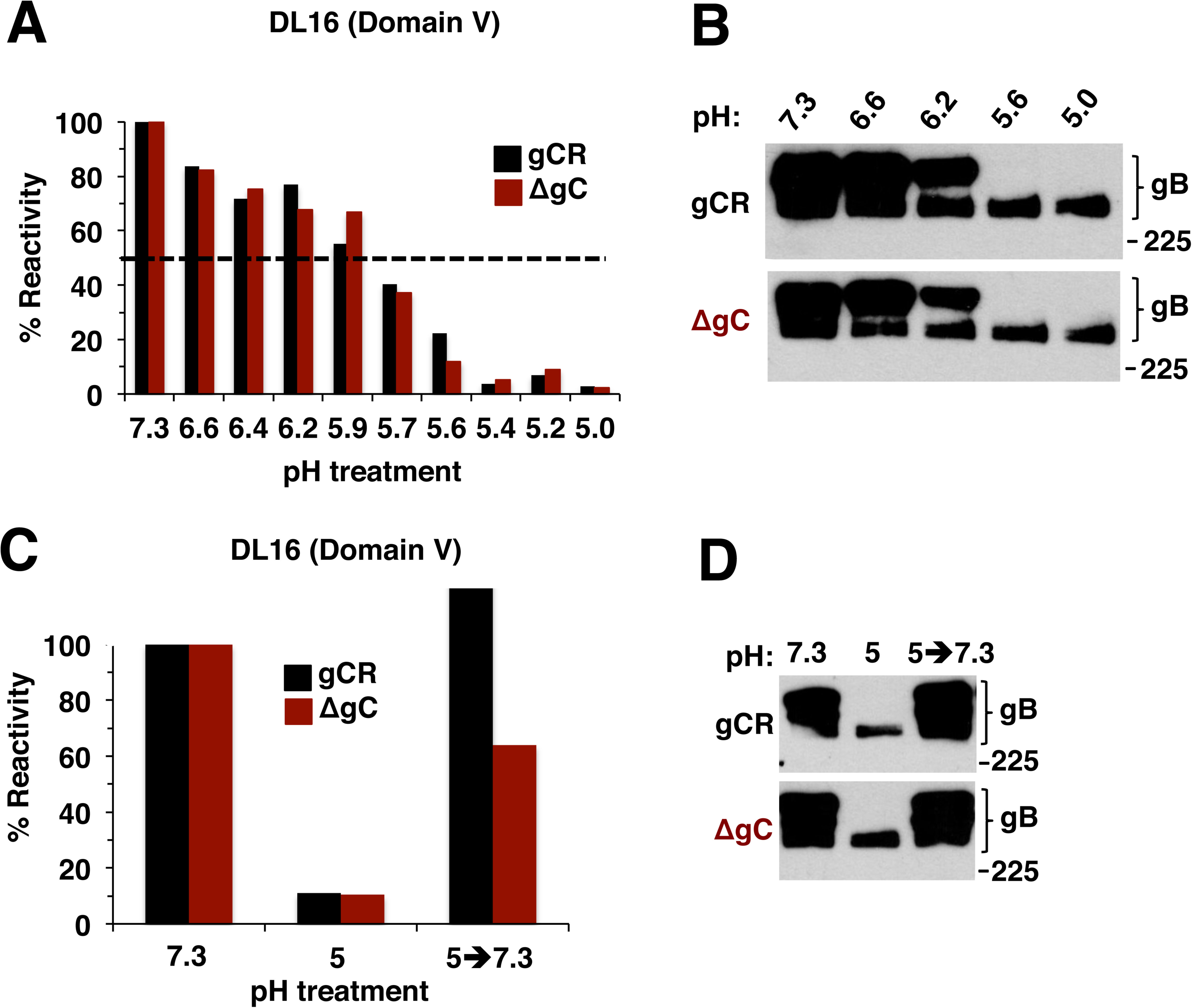
Low-pH-induced changes in gB oligomer are independent of gC as are their reversibility. (A) HSV-1 gCR or ΔgC was treated with a range of pHs for 10 min. Samples were directly blotted to nitrocellulose and probed at neutral pH with gB oligomer-specific MAb DL16. Reactivity was quantitated and the pH 7.3 sample was set as 100%. (B) gCR or ΔgC was treated with a range of pHs for 10 min. 1% SDS was added, and reactions were added to “native” PAGE sample buffer. Unheated samples were resolved by 8% SDS-PAGE and Western blot for gB. (C) gCR or ΔgC was treated with pH 5 for 10 min then blotted directly to nitrocellulose or first neutralized back to pH 7.3 for 10 min. Blots were probed at neutral pH with anti-gB MAb DL16. (D) gCR or ΔgC was treated with pH 5 for 10 min. One set was neutralized back to pH 7.3. 1% SDS was added and samples were processed as in panel B. Molecular size markers in kilodaltons are indicated at the right.

Since the changes in gB oligomeric conformation did not require gC, we expected that their reversibility would also be gC-independent. This was indeed the case. Acid-induced reduction of reactivity with the oligomer-specific MAb DL16 was partially reversible in both ΔgC and gCR viruses (Fig. 8C). Likewise, the susceptibility of low-pH treated gB oligomer to disruption by 1% SDS was also reversible, regardless of the presence of gC (Fig. 8D). Thus, the reversibility of low pH-triggered conformational changes in the gB oligomer is independent of gC.

## Discussion

The mechanisms underlying how herpesviruses traverse distinct cellular entry pathways is paramount to our understanding of these important pathogens. The experiments described here reveal a selective role for HSV-1 gC in low pH entry. HSV-1 gC confers an infectivity advantage in cells that support low pH entry of HSV-1, such as human keratinocytes. There is a lag in the exit of enveloped gC-negative particles from endocytic compartments, which may reflect a role for gC in optimal virus trafficking or low pH fusion. Low pH-triggered antigenic changes in gB domains I, II, and V are thought to be critical for fusion (16, 39, 40, 48–50); (Fig. 6 and S2). gC facilitates pH-induced gB conformational changes, increasing the pH of antigenic change by as much as 0.4 – 0.7 pH units. The reduced entry of gC-negative HSV may be explained by gC’s role in facilitating fusion-associated conformational changes in gB, which result in optimal penetration from an endocytic compartment. Importantly, in the absence of gC, HSV still uses a low-pH pathway to enter and infect cells, likely mediated in part by changes in gB that occur in the absence of gC, albeit at a lower pH. HSV-1 gC also binds to complement component C3b and inhibits complement-mediated immunity (51). When HSV particles are treated with soluble heparin in a cell-free assay, the UL16 tegument protein is rearranged in a gC-dependent manner (52). The role of gC in entry by an acidic endosomal pathway described here is independent of its role in attachment to cell surface heparan sulfate.

Viral envelope glycoprotein B (gB) is highly conserved among all subfamilies of the *Herpesviridae*. Current models of HSV-1 entry posit that 1) gC, and to a lesser extent gB, mediates viral attachment to host cell surface glycosaminoglycans; 2) gD binds to a cognate host cell receptor such as HVEM or nectin-1, resulting in pH-independent conformational change in gD; 3) This is thought to transmit a signal to gH/gL; and 4) culminate in the execution of membrane fusion by gB. Thus, gB, unlike other members of the class III fusion protein family, does not mediate fusion on its own. In addition to pH-triggered changes in gB domains I and V, we show here that acid-induced antigenic changes also occur in domain II (Fig. 6 and S2).

There is structure-based evidence that gB can exist in multiple conformations. The postfusion structure of gB is known, and distinct, membrane-associated non-postfusion forms, that may reflect the prefusion conformation, have also been resolved (53–55). MAbs that bind specifically to either pre-fusion or post-fusion gB have not been identified. Antibody binding to pre-fusion gB present in virions that are pretreated with low pH does not disappear completely; instead there are decreases in antibody reactivity. Low pH causes gB to assume a non-prefusion form, but is likely not sufficient to shift gB to the post-fusion form. In the absence of gC, the pH threshold for gB conformational changes, particularly in domain I and II is lower by 0.4-0.7 pH units (Fig 6). As a comparison, variants and mutants of influenza HA exhibit a ∼ 0.2-0.6 shift in the pH of both conformational change and fusion (56, 57). The H126 epitope in the fusion domain of gB might be particularly important for the pH-activation of fusion (18). In the absence of gC, a more acidic pH is required to trigger changes in the accessibility of the H126 epitope.

The cell tropism of herpesvirus entry and infection is influenced by subfamily-specific viral proteins. EBV gp42 is required for fusion and entry in B cells but not epithelial cells (58–62). The HCMV pentamer complex of envelope proteins is necessary for endosomal entry into epithelial and endothelial cells but not for pH-neutral entry into fibroblasts (25, 26, 63–65).

However, a detailed mechanism underlying EBV gp42 or the HCMV pentamer role in selection of entry pathway is not known. The alphaherpesvirus-specific protein gC is the first HSV envelope protein reported to selectively participate in endocytic entry. Here we suggest that gC is important for HSV-1 epithelial infection but less so for neuronal entry. The results are consistent with a mechanism whereby gC acts to ensure that gB undergoes optimal conformational change to mediate fusion with an appropriate endosomal compartment.

The results suggest a functional interaction between gC and gB. Direct interaction between gC and gB has not been detected by co-immunoprecipitation approaches at different pHs (data not shown). Low affinity or transient interactions may not be captured, or gC may exert an indirect effect on gB through another viral or host factor. Physical interactions between HSV-1 gB and gH have also been difficult to detect, despite demonstration of functional interactions (14, 15, 66). We propose a model whereby gC aids gB, and together with gD and gH/gL, allows rapid entry of HSV-1 into epithelial cells.

## Materials and Methods

### Cells and viruses

Human HaCaT epithelial keratinocytes and Vero cells were propagated in Dulbecco’s modified Eagle’s medium (DMEM; Thermo Fisher Scientific) supplemented with 10% fetal bovine serum (FBS; Atlanta Biologicals). Non-differentiated human SK-N-SH and IMR-32 neuroblastoma cells (ATCC) were propagated in Eagle’s minimal essential medium supplemented with 10% FBS, 1 mM sodium pyruvate, 0.1 mM nonessential amino acids, and Earle’s salts (Invitrogen). CHO-HVEM (M1A) cells (67) (provided by R. Eisenberg and G. Cohen, University of Pennsylvania) are stably transformed with the human HVEM gene and contain the *E. coli lacZ* gene under the control of the HSV-1 ICP4 gene promoter. CHO-HVEM cells were propagated in Ham’s F-12 nutrient mixture (Gibco/Life Technologies) supplemented with 10% FBS, 150 μg of puromycin (Sigma–Aldrich, St. Louis, MO, United States)/ml, and 250 μg of G418 sulfate (Thermo Fisher Scientific, Fair Lawn, NJ, United States)/ml. Cells were subcultured in non-selective medium prior to use in all experiments. Primary human epidermal keratinocytes (HEKa) (ATCC) up to passage 8 were maintained in dermal cell basal medium (ATCC) supplemented with keratinocyte growth kit (ATCC) and penicillin-streptomycin-amphotericin B solution (ATCC). CHOpgs745 cells (ATCC), which lack a gene required for heparan sulfate biosynthesis, were propagated in Ham’s F-12 nutrient mixture supplemented with 10% FBS. HSV-1 strain KOS and all viruses in this study were propagated and titered on Vero cells. HSV-1 (KOS) ΔgC2-3 virus or ΔgC is HSV-1 KOS in which most of the gC gene is deleted and replaced by *lacZ* (33); this virus is considered a gC-negative HSV-1. HSV-1 (KOS) gC2-3Rev virus or gCR is a recombinant in which HSV-1 (KOS) ΔgC2-3 was rescued by insertion of the wild-type gC gene (33). Both viruses were obtained from C. Brandt, University of Wisconsin, Madison. The viral genomes have not been sequenced. However, HSV-1 gCR and the parental KOS are indistinguishable in terms of specific infectivity, cell attachment, heparan sulfate binding, and infectivity. Their genomes are similar as measured by restriction enzyme analysis and Southern blotting (33). In addition, the phenotypes attributed to HSV-1 gCR in this study, are similar to those previously reported for HSV-1 wild type strain KOS (6, 7, 16, 18, 23, 68, 69). HSV-1 F-gE/GFP lacks the gE gene (70) (obtained from D. Johnson, Oregon Health Sciences University).

### Antibodies

Anti-gB mouse monoclonal antibodies H126 (domain I), H1359 (domain III), and H1817 (domain VI) were from Virusys. Anti-gB monoclonal antibodies DL16 (oligomer-specific; domain V), SS10 (domain IV), SS55 (domain I) (71), SS106 (domain V), and SS144 (domain V) (47) were provided by G. Cohen and R. Eisenberg, University of Pennsylvania. Anti-gB monoclonal antibodies H1838 and H1781 (domain II) were provided by L. Pereira, University of California, San Francisco (72).

### Immunofluorescence assay of HSV entry and infectivity

Equivalent amounts of HSV-1 ΔgC or gCR were added to cells grown on coverslips in 24-well plates in triplicate. Cells were incubated at 4°C for 1 hr then washed twice with cold PBS. Cultures were then shifted to 37°C for 6 hr then fixed with 100% ice-cold methanol. Primary antibody to HSV-1 ICP4 was then added, followed by Alexa Fluor-488-labeled secondary antibody. Nuclei were counterstained with 12.5 ng/ml DAPI. Approximately 500 cells per well were counted and scored for successful infection.

### HSV-1 plaque assay

HSV-1 titers were determined by limiting dilution plaque assay. At 18-20 hr p.i., cells were fixed with ice-cold methanol and acetone (2:1 ratio) for 20 min at −20°C and air-dried. Titers were determined by immunoperoxidase staining with rabbit polyclonal antibody to HSV, HR50 (Fitzgerald Industries, Concord, Mass.).

### Ectopic expression of HSV-1 gC

Lipofectamine 3000 (ThermoFisherScientific) was used to transfect cells with plasmids encoding gC (pSH140 obtained from G. Cohen and R. Eisenberg), gD (pPEP99; obtained from P. Spear), or empty vector. At 48 hr post-transfection, cells were infected with HSV-1, and infectivity was measured by plaque assay.

### SDS-PAGE and Western blotting

HSV-1 in Laemmli buffer with 200 mM dithiotheriol was boiled for 5 min. Proteins were separated by SDS-PAGE on Tris-glycine gels (Thermo Fisher Scientific). For protein staining, gels were then fixed and stained with 0.025% Coomasie brilliant blue (J.T. Baker Chemical Co., Philipsburg, NJ), 40% methanol (Baker Chemical), and 10% glacial acetic acid (Baker Chemical), followed by destaining with 30% methanol and 7% glacial acetic acid (69). Gels were dried and imaged with a Gel Doc XR imager (Bio-Rad, Hercules, CA). For Western blotting, following transfer to nitrocellulose, membranes were blocked and incubated with HSV polyclonal antibodies R68 (α-gB), R47 (α-gC) (73), R2 (α-gD) (74), R137 (α-gH), anti-gD monoclonal antibody DL6 (75), which were gifts from Gary Cohen and Roselyn Eisenberg; HSV-1 gE monoclonal antibody H1A054-100 (Virusys); or monoclonal antibody H1A021 to VP5 (Santa Cruz Biotechnology, Dallas, TX). After incubation with horseradish peroxidase-conjugated secondary antibodies, enhanced chemiluminescent substrate (Pierce) was added, and membranes were exposed to X-ray film (Kodak).

### Effect of ammonium chloride on HSV entry and infectivity

Cells grown in 24-well plates were treated with medium containing ammonium chloride for 1 hr at 37°C. Virus was added in the continued presence of agent for 6 h. Medium was then removed and replaced with complete DMEM. At 16 h p.i., plaque assay was performed to measure infectivity. CHO-HVEM cells grown in 96-well plates were treated with medium containing ammonium chloride for 20 min at 37°C. Virus was added (MOI of 5) in the continued presence of agent, and beta-galactosidase activity of cell lysates was measured at 6 h p.i.

### Attachment of HSV-1 to the cell surface

Cells grown in 96-well plates were pre-chilled in carbonate-free, serum-free medium supplemented with 20 mM HEPES and 0.2% bovine serum albumin (binding medium) at 4°C on ice for 20 min. Approximately 10^6^ particles of HSV-1 ΔgC or gCR in binding medium were added to cells at 4°C on ice for 1 hr. Cultures were washed twice with ice-cold PBS. Cells were trypsinized, and cell-associated HSV-1 DNA was isolated with the QIAamp DNA Blood Mini Kit (Qiagen) according to the manufacturer’s instructions. Attached HSV-1 was quantitated by qPCR as previously described (11, 76).

### Intracellular tracking of enveloped, infectious HSV

HSV-1 ΔgC or gCR was added to confluent cell monolayers (MOI of 8) on ice at 4°C for 2 hr. Cultures were washed with PBS and shifted to 37°C. At the indicated times p.i., extracellular virus was inactivated by adding sodium citrate buffer (pH 3.0) for 1 min at 37°C (7). Monolayers were immediately put on ice and washed with cold PBS. One milliliter of Ham’s F12 medium with 20 mM HEPES and 1% FBS was added, and cells were lysed by two cycles of freezing and thawing. Titers of lysates were determined on Vero cells. Each data point is the mean of triplicate samples.

### Dot blot analysis of HSV-1 gB

Extracellular HSV preparations were diluted in serum-free, bicarbonate-free DMEM with 0.2% bovine serum albumin (BSA) and 5 mM each of HEPES, MES, and sodium succinate. Virions were adjusted with HCl to achieve a range of different pHs from 5.0 to 7.3. Samples were incubated at 37°C for 10 min, and then either blotted directly to nitrocellulose membrane using a Minifold dot blot system (Whatman), or were first neutralized by addition of pretitrated amounts of 0.05 N NaOH (77). Virus-dotted nitrocellulose membranes were blocked, and then incubated with antibodies to gB at neutral pH. After incubation with horseradish peroxidase-conjugated secondary antibodies, enhanced chemiluminescent substrate (Thermo Fisher Scientific) was added, and blots were exposed to X-ray film (Genesee Scientific). Densitometry was performed with ImageJ.

### Analysis of gB oligomeric structure by PAGE

Extracellular preparations of HSV were diluted in the same medium used for dot blot analysis. Virus samples were adjusted to the indicated pHs with pretitrated amounts of 0.05 N HCl for 10 min at 37°C. To test for reversibility of conformational change, samples were then neutralized by addition of pretitrated amounts of 0.05 N NaOH for 10 min at 37°C. 1% SDS was then added. Laemmli sample buffer containing 0.2% sodium dodecyl sulfate (SDS) with no reducing agent (78) was added. Samples were not heated, and proteins were resolved by PAGE. After transfer to nitrocellulose, membranes were blocked and incubated with gB monoclonal antibody H1359. After incubation with horseradish peroxidase-conjugated secondary antibody, enhanced chemiluminescent substrate (Thermo Fisher) was added and membranes were exposed to X-ray film (Genesee Scientific).

## Acknowledgments

We thank Ryan Manglona, Erik Walker, George Wudiri, and Youki Yamasaki for early contributions to this work. We are grateful to Curtis Brandt, Gary Cohen, Roselyn Eisenberg, David Johnson, Lenore Pereira, and Patricia Spear for providing reagents. This study was supported by National Institutes of Health (NIH) grant R01 AI119159 (A.V.N) and NIH Training Grant T32 GM008336 (T.K., D.J.W., and K.A.G.).

---

**Fig. S1.**
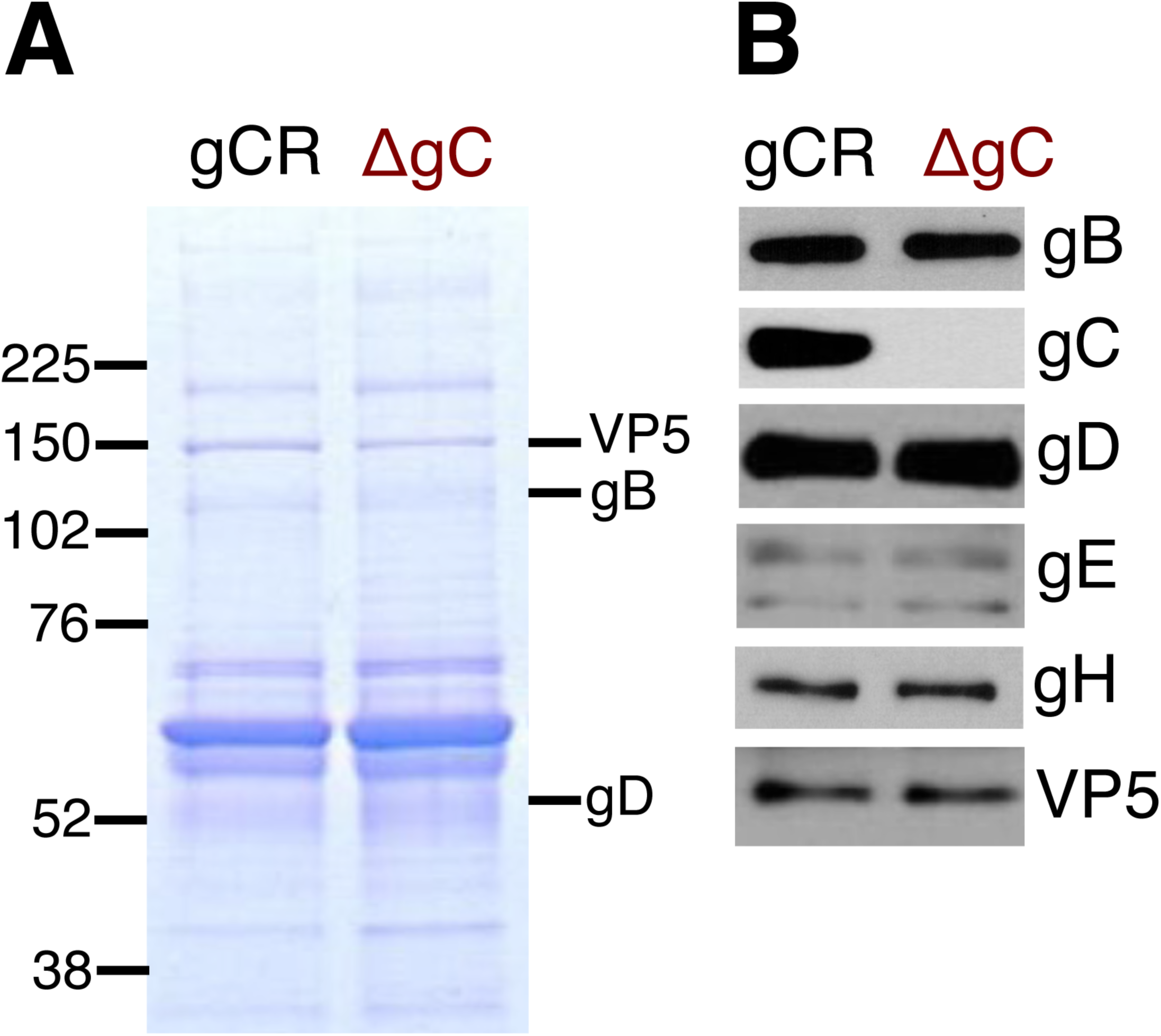
Protein composition of gC-negative HSV-1 ΔgC. Equivalent VP5 units of HSV-1 gCR or ΔgC in Laemmli buffer were separated by SDS-PAGE followed by (A) protein staining with Coomassie blue or (B) Western blotting for the indicated viral proteins. Molecular weight standards in kilodaltons are indicated to the left.

**Fig. S2.**
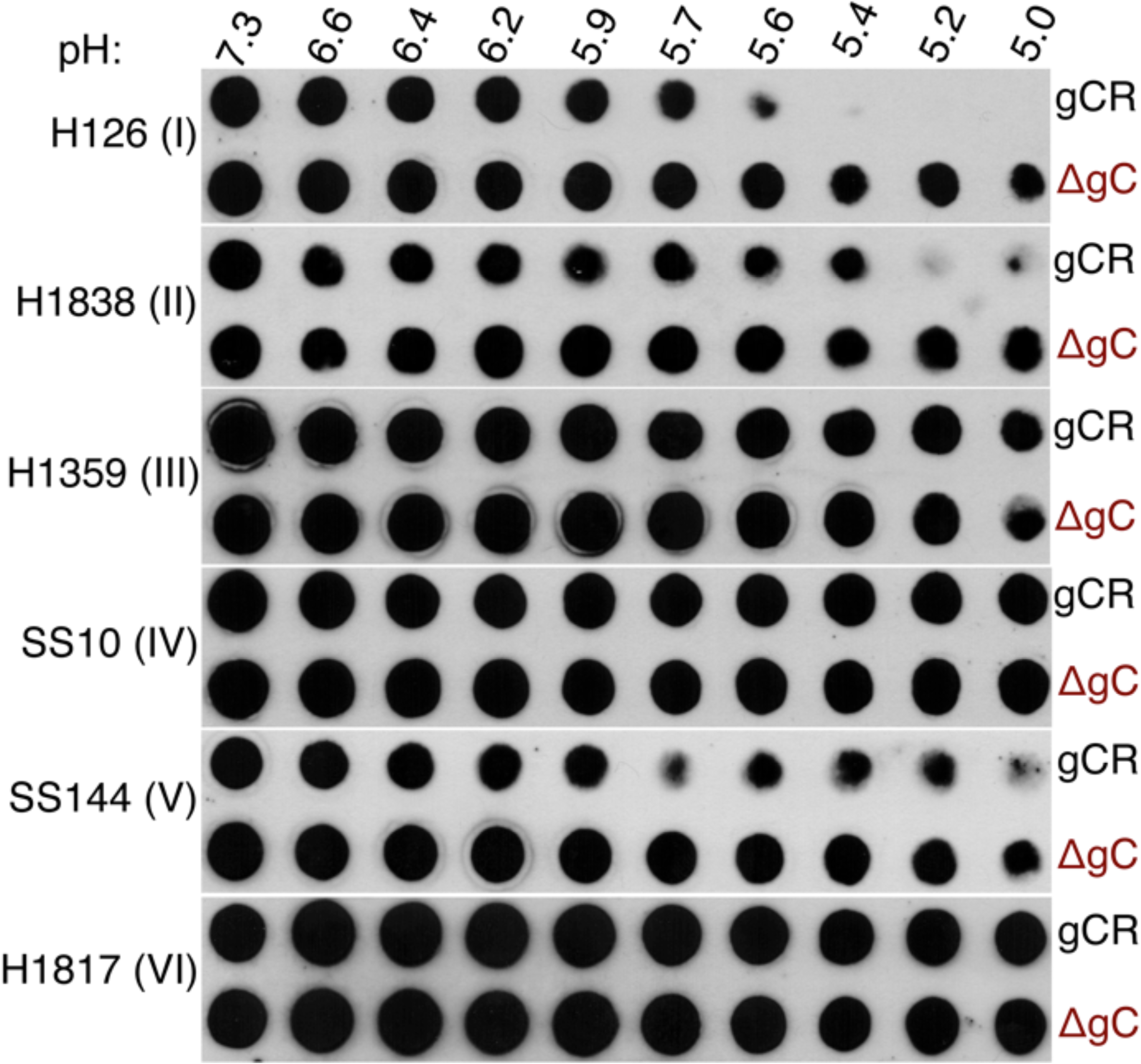
HSV-1 gC facilitates acid-induced conformational changes in the fusion protein gB. Extracellular preparations of HSV-1 gCR or ΔgC (∼ 10^7^ genome copy numbers) were treated with pHs ranging from 7.3 to 5.0 and blotted directly to nitrocellulose. Blots were probed with the indicated gB MAbs H126, H1838, SS144, or H1359 at neutral pH followed by HRP-conjugated anti-mouse secondary antibody. The antibody name is shown at the left and the gB domain to which each MAb is directed is indicated in parenthesis. These are individual examples of experiments that were quantitated and averaged together with multiple similar independent determinations. Summarized quantitative results are depicted in Figure 6.

**Fig. S3.**
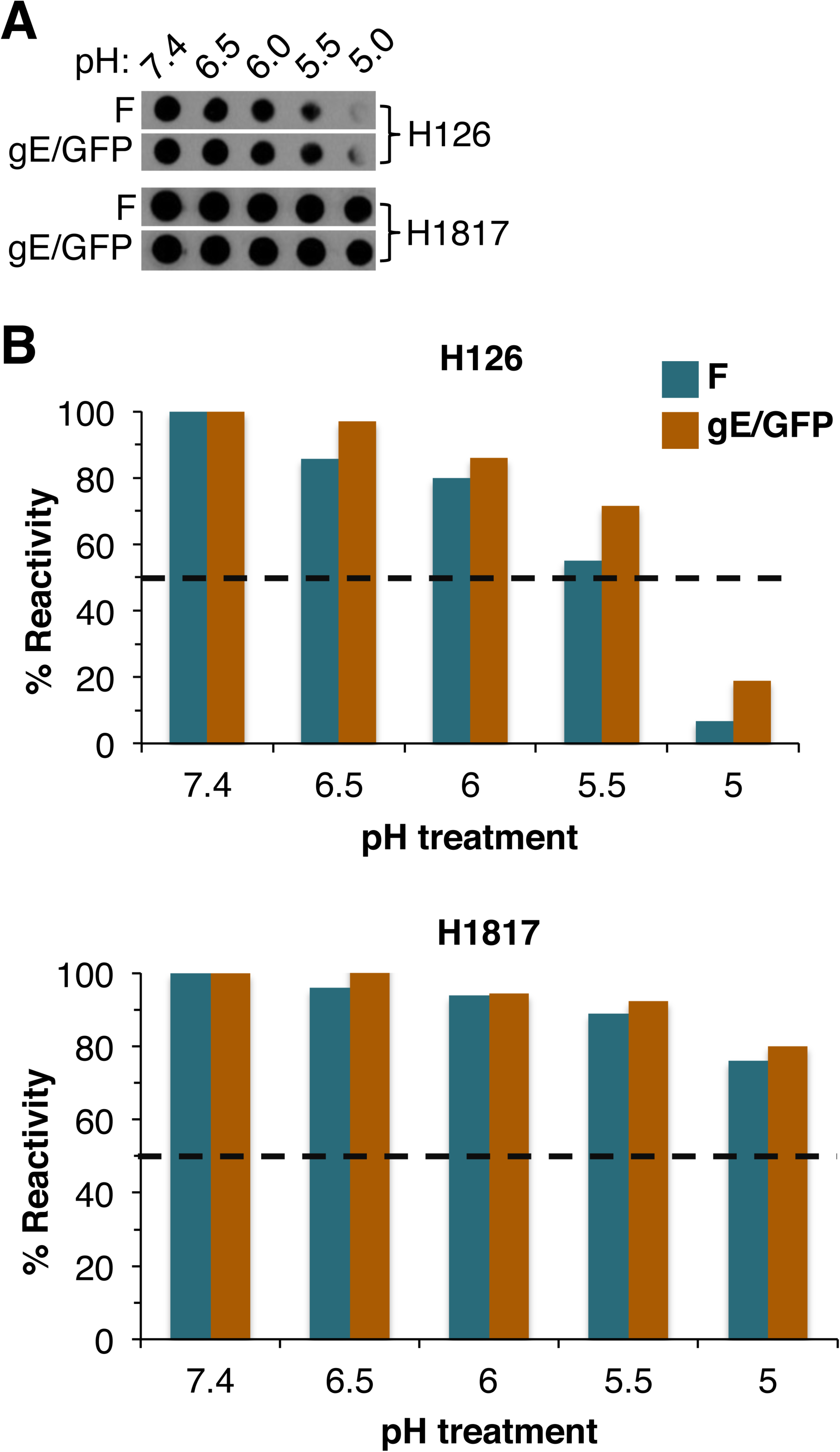
HSV-1 gE does not influence acid-induced conformational change in the H126 epitope of gB. (A) HSV-1 strain wild type F or its derivative gE-GFP were treated with indicated pHs then directly blotted to nitrocellulose membrane. Blot was probed with representative gB MAbs H126 or H1817 at neutral pH. (B) Antibody reactivity was quantitated and treatment with pH 7.4 was set as 100%.

**Fig. S4.**
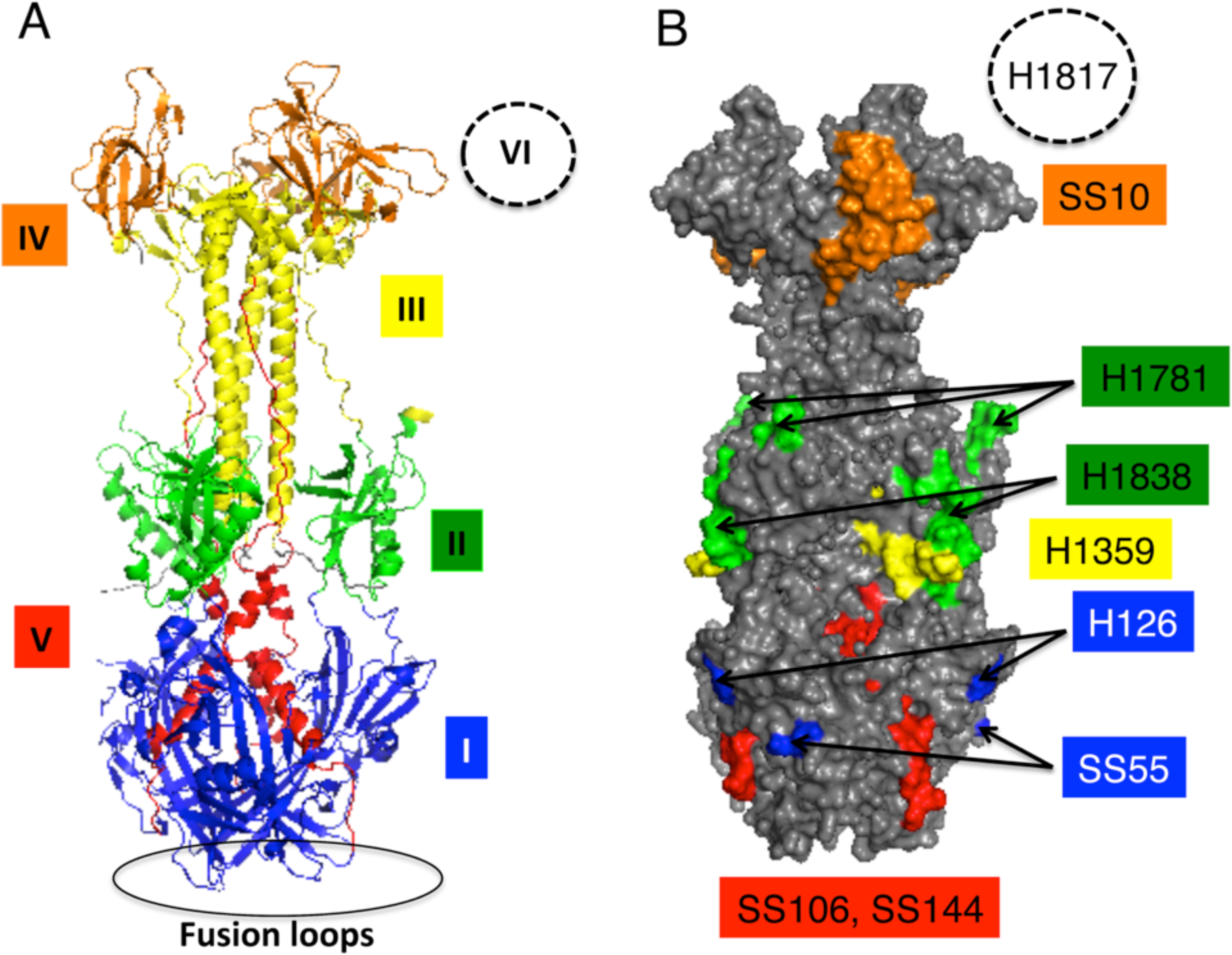
Domain structure of HSV-1 gB and location of MAb epitopes. (A) gB ectodomain trimer representing a post-fusion conformation. (B) Location of monoclonal antibody-binding sites. Monoclonal antibody-resistant mutations in Domain I, which contains bipartite hydrophobic fusion loops, map to amino acid residue 303 for H126 and residues 203, 335, and 199 for SS55 (79, 80). The MAb H1781 epitope in Domain II maps to 454-473, and H1838 maps to 391-410 (47). The H1359 epitope in Domain III maps to 487–505 (72). SS10 in Domain IV maps to 640–670 (47). SS106 and SS144 to Domain V both bind to 697–725 (53). The MAb H1817 epitope in Domain VI (not resolved in the structure) maps to 31–43 (47).

## References

1. Nicola AV. 2016. Herpesvirus Entry into Host Cells Mediated by Endosomal Low pH. Traffic 17:965–975.

2. Sathiyamoorthy K, Chen J, Longnecker R, Jardetzky TS. 2017. The COMPLEXity in herpesvirus entry. Curr Opin Virol 24:97–104.

3. Krummenacher C, Carfi A, Eisenberg RJ, Cohen GH. 2013. Entry of herpesviruses into cells: the enigma variations. Adv Exp Med Biol 790:178–195.

4. Weed DJ, Nicola AV. 2017. Herpes simplex virus Membrane Fusion. Adv Anat Embryol Cell Biol 223:29–47.

5. Campadelli-Fiume G, Menotti L, Avitabile E, Gianni T. 2012. Viral and cellular contributions to herpes simplex virus entry into the cell. Current opinion in virology 2:28–36.

6. Nicola AV, Hou J, Major EO, Straus SE. 2005. Herpes simplex virus type 1 enters human epidermal keratinocytes, but not neurons, via a pH-dependent endocytic pathway. Journal of virology 79:7609–7616.

7. Nicola AV, McEvoy AM, Straus SE. 2003. Roles for endocytosis and low pH in herpes simplex virus entry into HeLa and Chinese hamster ovary cells. Journal of virology 77:5324–5332.

8. Geraghty RJ, Krummenacher C, Cohen GH, Eisenberg RJ, Spear PG. 1998. Entry of alphaherpesviruses mediated by poliovirus receptor-related protein 1 and poliovirus receptor. Science 280:1618–1620.

9. Spear PG, Eisenberg RJ, Cohen GH. 2000. Three classes of cell surface receptors for alphaherpesvirus entry. Virology 275:1–8.

10. Campadelli-Fiume G, Cocchi F, Menotti L, Lopez M. 2000. The novel receptors that mediate the entry of herpes simplex viruses and animal alphaherpesviruses into cells. Rev Med Virol 10:305–319.

11. Komala Sari T, Pritchard SM, Cunha CW, Wudiri GA, Laws EI, Aguilar HC, Taus NS, Nicola AV. 2013. Contributions of herpes simplex virus 1 envelope proteins to entry by endocytosis. Journal of virology 87:13922–13926.

12. Baquero E, Albertini AA, Gaudin Y. 2015. Recent mechanistic and structural insights on class III viral fusion glycoproteins. Curr Opin Struct Biol 33:52–60.

13. Chowdary TK, Cairns TM, Atanasiu D, Cohen GH, Eisenberg RJ, Heldwein EE. 2010. Crystal structure of the conserved herpesvirus fusion regulator complex gH-gL. Nat Struct Mol Biol 17:882–888.

14. Atanasiu D, Whitbeck JC, Cairns TM, Reilly B, Cohen GH, Eisenberg RJ. 2007. Bimolecular complementation reveals that glycoproteins gB and gH/gL of herpes simplex virus interact with each other during cell fusion. Proc Natl Acad Sci U S A 104:18718–18723.

15. Avitabile E, Forghieri C, Campadelli-Fiume G. 2007. Complexes between herpes simplex virus glycoproteins gD, gB, and gH detected in cells by complementation of split enhanced green fluorescent protein. J Virol 81:11532–11537.

16. Dollery SJ, Delboy MG, Nicola AV. 2010. Low pH-induced conformational change in herpes simplex virus glycoprotein B. Journal of virology 84:3759–3766.

17. Gallagher JR, Atanasiu D, Saw WT, Paradisgarten MJ, Whitbeck JC, Eisenberg RJ, Cohen GH. 2014. Functional fluorescent protein insertions in herpes simplex virus gB report on gB conformation before and after execution of membrane fusion. PLoS Pathog 10:e1004373.

18. Weed DJ, Pritchard SM, Gonzalez F, Aguilar HC, Nicola AV. 2017. Mildly Acidic pH Triggers an Irreversible Conformational Change in the Fusion Domain of Herpes Simplex Virus 1 Glycoprotein B and Inactivation of Viral Entry. J Virol 91.

19. Roller DG, Dollery SJ, Doyle JL, Nicola AV. 2008. Structure-function analysis of herpes simplex virus glycoprotein B with fusion-from-without activity. Virology 382:207–216.

20. Herold BC, WuDunn D, Soltys N, Spear PG. 1991. Glycoprotein C of herpes simplex virus type 1 plays a principal role in the adsorption of virus to cells and in infectivity. Journal of virology 65:1090–1098.

21. WuDunn D, Spear PG. 1989. Initial interaction of herpes simplex virus with cells is binding to heparan sulfate. J Virol 63:52–58.

22. Shukla D, Spear PG. 2001. Herpesviruses and heparan sulfate: an intimate relationship in aid of viral entry. J Clin Invest 108:503–510.

23. Nicola AV, Straus SE. 2004. Cellular and viral requirements for rapid endocytic entry of herpes simplex virus. Journal of virology 78:7508–7517.

24. Akula SM, Naranatt PP, Walia NS, Wang FZ, Fegley B, Chandran B. 2003. Kaposi’s sarcoma-associated herpesvirus (human herpesvirus 8) infection of human fibroblast cells occurs through endocytosis. J Virol 77:7978–7990.

25. Ryckman BJ, Jarvis MA, Drummond DD, Nelson JA, Johnson DC. 2006. Human cytomegalovirus entry into epithelial and endothelial cells depends on genes UL128 to UL150 and occurs by endocytosis and low-pH fusion. Journal of virology 80:710–722.

26. Wang D, Yu QC, Schroer J, Murphy E, Shenk T. 2007. Human cytomegalovirus uses two distinct pathways to enter retinal pigmented epithelial cells. Proc Natl Acad Sci U S A 104:20037–20042.

27. Gianni T, Campadelli-Fiume G, Menotti L. 2004. Entry of herpes simplex virus mediated by chimeric forms of nectin1 retargeted to endosomes or to lipid rafts occurs through acidic endosomes. Journal of virology 78:12268–12276.

28. Finnen RL, Mizokami KR, Banfield BW, Cai GY, Simpson SA, Pizer LI, Levin MJ. 2006. Postentry events are responsible for restriction of productive varicella-zoster virus infection in Chinese hamster ovary cells. J Virol 80:10325–10334.

29. Cai WH, Gu B, Person S. 1988. Role of glycoprotein B of herpes simplex virus type 1 in viral entry and cell fusion. J Virol 62:2596–2604.

30. Ligas MW, Johnson DC. 1988. A herpes simplex virus mutant in which glycoprotein D sequences are replaced by beta-galactosidase sequences binds to but is unable to penetrate into cells. Journal of Virology 62:1486–1494.

31. Forrester A, Farrell H, Wilkinson G, Kaye J, Davis-Poynter N, Minson T. 1992. Construction and properties of a mutant of herpes simplex virus type 1 with glycoprotein H coding sequences deleted. Journal of virology 66:341–348.

32. Roop C, Hutchinson L, Johnson DC. 1993. A mutant herpes simplex virus type 1 unable to express glycoprotein L cannot enter cells, and its particles lack glycoprotein H. Journal of virology 67:2285–2297.

33. Herold BC, Visalli RJ, Susmarski N, Brandt CR, Spear PG. 1994. Glycoprotein C-independent binding of herpes simplex virus to cells requires cell surface heparan sulphate and glycoprotein B. The Journal of general virology 75 (Pt 6):1211–1222.

34. Campadelli-Fiume G, Stirpe D, Boscaro A, Avitabile E, Foa-Tomasi L, Barker D, Roizman B. 1990. Glycoprotein C-dependent attachment of herpes simplex virus to susceptible cells leading to productive infection. Virology 178:213–222.

35. Koyama AH, Uchida T. 1987. The mode of entry of herpes simplex virus type 1 into Vero cells. Microbiology and immunology 31:123–130.

36. Wittels M, Spear PG. 1991. Penetration of cells by herpes simplex virus does not require a low pH-dependent endocytic pathway. Virus research 18:271–290.

37. Johnson RM, Spear PG. 1989. Herpes simplex virus glycoprotein D mediates interference with herpes simplex virus infection. J Virol 63:819–827.

38. Holland TC, Homa FL, Marlin SD, Levine M, Glorioso J. 1984. Herpes simplex virus type 1 glycoprotein C-negative mutants exhibit multiple phenotypes, including secretion of truncated glycoproteins. J Virol 52:566–574.

39. Siekavizza-Robles CR, Dollery SJ, Nicola AV. 2010. Reversible conformational change in herpes simplex virus glycoprotein B with fusion-from-without activity is triggered by mildly acidic pH. Virology journal 7:352.

40. Dollery SJ, Wright CC, Johnson DC, Nicola AV. 2011. Low-pH-dependent changes in the conformation and oligomeric state of the prefusion form of herpes simplex virus glycoprotein B are separable from fusion activity. Journal of virology 85:9964–9973.

41. Hannah BP, Heldwein EE, Bender FC, Cohen GH, Eisenberg RJ. 2007. Mutational evidence of internal fusion loops in herpes simplex virus glycoprotein B. J Virol 81:4858–4865.

42. Hannah BP, Cairns TM, Bender FC, Whitbeck JC, Lou H, Eisenberg RJ, Cohen GH. 2009. Herpes simplex virus glycoprotein B associates with target membranes via its fusion loops. J Virol 83:6825–6836.

43. Clement C, Tiwari V, Scanlan PM, Valyi-Nagy T, Yue BY, Shukla D. 2006. A novel role for phagocytosis-like uptake in herpes simplex virus entry. J Cell Biol 174:1009–1021.

44. Zhou J, Blissard GW. 2006. Mapping the conformational epitope of a neutralizing antibody (AcV1) directed against the AcMNPV GP64 protein. Virology 352:427–437.

45. Gaudin Y, Tuffereau C, Segretain D, Knossow M, Flamand A. 1991. Reversible conformational changes and fusion activity of rabies virus glycoprotein. J Virol 65:4853–4859.

46. Gaudin Y. 2000. Reversibility in fusion protein conformational changes. The intriguing case of rhabdovirus-induced membrane fusion. Subcell Biochem 34:379–408.

47. Bender FC, Samanta M, Heldwein EE, de Leon MP, Bilman E, Lou H, Whitbeck JC, Eisenberg RJ, Cohen GH. 2007. Antigenic and mutational analyses of herpes simplex virus glycoprotein B reveal four functional regions. J Virol 81:3827–3841.

48. Cairns TM, Whitbeck JC, Lou H, Heldwein EE, Chowdary TK, Eisenberg RJ, Cohen GH. 2011. Capturing the herpes simplex virus core fusion complex (gB-gH/gL) in an acidic environment. Journal of virology 85:6175–6184.

49. Stampfer SD, Lou H, Cohen GH, Eisenberg RJ, Heldwein EE. 2010. Structural basis of local, pH-dependent conformational changes in glycoprotein B from herpes simplex virus type 1. J Virol 84:12924–12933.

50. Muggeridge MI. 2012. Glycoprotein B of herpes simplex virus 2 has more than one intracellular conformation and is altered by low pH. J Virol 86:6444–6456.

51. Friedman HM, Cohen GH, Eisenberg RJ, Seidel CA, Cines DB. 1984. Glycoprotein C of herpes simplex virus 1 acts as a receptor for the C3b complement component on infected cells. Nature 309:633–635.

52. Meckes DG, Jr., Wills JW. 2008. Structural rearrangement within an enveloped virus upon binding to the host cell. J Virol 82:10429–10435.

53. Heldwein EE, Lou H, Bender FC, Cohen GH, Eisenberg RJ, Harrison SC. 2006. Crystal structure of glycoprotein B from herpes simplex virus 1. Science 313:217–220.

54. Fontana J, Atanasiu D, Saw WT, Gallagher JR, Cox RG, Whitbeck JC, Brown LM, Eisenberg RJ, Cohen GH. 2017. The Fusion Loops of the Initial Prefusion Conformation of Herpes Simplex Virus 1 Fusion Protein Point Toward the Membrane. MBio 8.

55. Zeev-Ben-Mordehai T, Vasishtan D, Hernandez Duran A, Vollmer B, White P, Prasad Pandurangan A, Siebert CA, Topf M, Grunewald K. 2016. Two distinct trimeric conformations of natively membrane-anchored full-length herpes simplex virus 1 glycoprotein B. Proc Natl Acad Sci U S A 113:4176–4181.

56. Daniels RS, Downie JC, Hay AJ, Knossow M, Skehel JJ, Wang ML, Wiley DC. 1985. Fusion mutants of the influenza virus hemagglutinin glycoprotein. Cell 40:431–439.

57. Doms RW, Gething MJ, Henneberry J, White J, Helenius A. 1986. Variant influenza virus hemagglutinin that induces fusion at elevated pH. J Virol 57:603–613.

58. Wang X, Hutt-Fletcher LM. 1998. Epstein-Barr virus lacking glycoprotein gp42 can bind to B cells but is not able to infect. Journal of virology 72:158–163.

59. Wang X, Kenyon WJ, Li Q, Mullberg J, Hutt-Fletcher LM. 1998. Epstein-Barr virus uses different complexes of glycoproteins gH and gL to infect B lymphocytes and epithelial cells. Journal of virology 72:5552–5558.

60. Miller N, Hutt-Fletcher LM. 1992. Epstein-Barr virus enters B cells and epithelial cells by different routes. Journal of virology 66:3409–3414.

61. Nemerow GR, Cooper NR. 1984. Early events in the infection of human B lymphocytes by Epstein-Barr virus: the internalization process. Virology 132:186–198.

62. Connolly SA, Jackson JO, Jardetzky TS, Longnecker R. 2011. Fusing structure and function: a structural view of the herpesvirus entry machinery. Nature reviews Microbiology 9:369–381.

63. Wang D, Shenk T. 2005. Human cytomegalovirus UL131 open reading frame is required for epithelial cell tropism. Journal of Virology 79:10330–10338.

64. Wang D, Shenk T. 2005. Human cytomegalovirus virion protein complex required for epithelial and endothelial cell tropism. Proceedings of the National Academy of Sciences of the United States of America 102:18153–18158.

65. Vanarsdall AL, Johnson DC. 2012. Human cytomegalovirus entry into cells. Current opinion in virology 2:37–42.

66. Vanarsdall AL, Howard PW, Wisner TW, Johnson DC. 2016. Human Cytomegalovirus gH/gL Forms a Stable Complex with the Fusion Protein gB in Virions. PLoS Pathog 12:e1005564.

67. Montgomery RI, Warner MS, Lum BJ, Spear PG. 1996. Herpes simplex virus-1 entry into cells mediated by a novel member of the TNF/NGF receptor family. Cell 87:427–436.

68. Delboy MG, Siekavizza-Robles CR, Nicola AV. 2010. Herpes simplex virus tegument ICP0 is capsid associated, and its E3 ubiquitin ligase domain is important for incorporation into virions. Journal of virology 84:1637–1640.

69. Wudiri GA, Schneider SM, Nicola AV. 2017. Herpes Simplex Virus 1 Envelope Cholesterol Facilitates Membrane Fusion. Front Microbiol 8:2383.

70. Farnsworth A, Goldsmith K, Johnson DC. 2003. Herpes simplex virus glycoproteins gD and gE/gI serve essential but redundant functions during acquisition of the virion envelope in the cytoplasm. Journal of virology 77:8481–8494.

71. Bender FC, Whitbeck JC, Lou H, Cohen GH, Eisenberg RJ. 2005. Herpes simplex virus glycoprotein B binds to cell surfaces independently of heparan sulfate and blocks virus entry. J Virol 79:11588–11597.

72. Pereira L, Ali M, Kousoulas K, Huo B, Banks T. 1989. Domain structure of herpes simplex virus 1 glycoprotein B: neutralizing epitopes map in regions of continuous and discontinuous residues. Virology 172:11–24.

73. Eisenberg RJ, Ponce de Leon M, Friedman HM, Fries LF, Frank MM, Hastings JC, Cohen GH. 1987. Complement component C3b binds directly to purified glycoprotein C of herpes simplex virus types 1 and 2. Microb Pathog 3:423–435.

74. Isola VJ, Eisenberg RJ, Siebert GR, Heilman CJ, Wilcox WC, Cohen GH. 1989. Fine mapping of antigenic site II of herpes simplex virus glycoprotein D. J Virol 63:2325–2334.

75. Eisenberg RJ, Long D, Ponce de Leon M, Matthews JT, Spear PG, Gibson MG, Lasky LA, Berman P, Golub E, Cohen GH. 1985. Localization of epitopes of herpes simplex virus type 1 glycoprotein D. J Virol 53:634–644.

76. Walker EB, Pritchard SM, Cunha CW, Aguilar HC, Nicola AV. 2015. Polyethylene glycol-mediated fusion of herpes simplex type 1 virions with the plasma membrane of cells that support endocytic entry. Virol J 12:190.

77. Sari TK, Gianopulos KA, Nicola AV. 2020. Conformational Change in Herpes Simplex Virus Entry Glycoproteins Detected by Dot Blot. 2060:319–326.

78. Cohen GH, Isola VJ, Kuhns J, Berman PW, Eisenberg RJ. 1986. Localization of discontinuous epitopes of herpes simplex virus glycoprotein D: use of a nondenaturing (“native” gel) system of polyacrylamide gel electrophoresis coupled with Western blotting. J Virol 60:157–166.

79. Kousoulas KG, Huo B, Pereira L. 1988. Antibody-resistant mutations in cross-reactive and type-specific epitopes of herpes simplex virus 1 glycoprotein B map in separate domains. Virology 166:423–431.

80. Cairns TM, Fontana J, Huang ZY, Whitbeck JC, Atanasiu D, Rao S, Shelly SS, Lou H, Ponce de Leon M, Steven AC, Eisenberg RJ, Cohen GH. 2014. Mechanism of neutralization of herpes simplex virus by antibodies directed at the fusion domain of glycoprotein B. J Virol 88:2677–2689.

